# Achiasmatic meiosis underlies propagation of clonal genome and B chromosomes in hexaploid *Carassius gibelio* females but not males

**DOI:** 10.64898/2026.05.30.728974

**Authors:** Dmitrij Dedukh, Heorhii Zolotarov, Karolina Komashchuk, Manfred Schartl, Josef Wanzenböck, Zuzana Majtanova, Yukiko Imai, Vladimir Trifonov, Karel Janko, Dunja K Lamatsch

**Author notes:** Corresponding author –.

## Abstract

Sexual reproduction in eukaryotes relies on meiosis, recombination, and fertilization, yet hybridization can lead to transitions to asexuality. Asexual vertebrate hybrids require modified gametogenesis to produce unreduced gametes, but the underlying mechanisms remain poorly understood in various hybrid complexes. Here, we investigated the *Carassius gibelio* complex, which combines clonal genome propagation to the offspring along with the transmission of B chromosomes. We analyzed meiotic progression and gametogenesis in sexual tetraploid and asexual hexaploid lineages, focusing on sex-specific differences. Cytological analyses of synaptonemal complexes and diplotene chromosomes revealed that both sexes in hexaploid lineages undergo achiasmatic meiosis, characterized by the absence of homologous pairing, recombination, and chiasmata. Chromosomes persist as univalents throughout meiotic prophase. Despite this shared meiotic program, outcomes differ between sexes: females complete oogenesis and produce unreduced eggs, whereas males exhibit disrupted spermatogenesis and reduced fertility. Females bypass the reductional division, enabling clonal gamete formation, while males fail to segregate univalents properly. Furthermore, B chromosomes were detected in both mitosis and meiosis of hexaploid males and females, but not in sexual lineages. B chromosomes were consistently present and varied in number among individuals. B chromosomes varied in number and consistently formed univalents during meiosis, similar to other chromosomes.

**Significance statement:** Meiosis normally depends on chromosome pairing and recombination, yet asexual vertebrates can bypass these processes. We demonstrate that asexual hexaploid *Carassius gibelio* reproduces through achiasmatic meiosis, in which chromosomes fail to pair and recombine. While females successfully produce unreduced clonal eggs, males display reduced fertility, revealing striking sex-specific differences in the ability to overcome meiotic dysfunction. We further show that B chromosomes persist as meiotic univalents and are stably inherited.

## Introduction

Sexual reproduction is a defining feature of eukaryotes and relies on meiosis, crossing over, fertilization, and the development of new individuals [1,2]. Despite the universality of sexual reproduction and its ancestral state in all eukaryotes, asexual reproduction also evolved as a spin-off at the tips of many phylogenetic lineages [3–5]. Although asexuals are found in many taxa, they remain relatively rare, and their overall frequency has not been systematically quantified, among others because the term “asexuality” is loosely defined [3,5,6]. In one of the strict meaning, the organisms called as asexual transmit their genomes clonally to their offspring without recombination, producing genetically identical progeny [3,5,7]. However, clonal transmission is not exclusive to wholly asexual organisms, as even within otherwise sexually reproducing species, certain genomic elements are known to deviate from Mendelian inheritance and propagate outside the rules of canonical meiosis and fertilization. Supernumerary elements such as B chromosomes and germline-restricted chromosomes represent prominent examples of such non-Mendelian inheritance, as they are transmitted clonally or through drive mechanisms independently of the rest of the genome [8,9]. Although the mechanisms of clonal transmission of whole genomes or individual chromosomes vary widely, they generally do not conform to canonical premeiotic, meiotic, or fertilization processes specific for sexual reproduction.

Hybridization often affects canonical reproductive pathways, leading to reduced fertility or sterility in hybrids [10,11]. These effects typically arise from improper chromosome pairing and recombination during meiosis, leading to postzygotic reproductive isolation. In taxa with heteromorphic sex chromosomes, such disruptions disproportionately affect the heterogametic sex, consistent with Haldane’s rule [10–13]. Despite these barriers, hybridization between sufficiently related species may give rise to diverse forms of asexuality [3–5]. Such asexual hybrids generally emerge at a certain level of genetic divergence and establish clonal or hemiclonal lineages through modified gametogenesis [3,5–7,14]. In hybrid vertebrates, alterations in gamete production, whether premeiotic or meiotic, can overcome hybrid sterility and enable the production of clonal or hemiclonal gametes [3,5,7,14]. Further alterations of fertilization may further stabilize clonal reproduction without introgression from parental genomes [9,15,16]. These processes have given rise to diverse reproductive modes, including parthenogenesis, gynogenesis, and hybridogenesis, which have independently evolved across multiple taxa [3,5,7].

In asexual organisms, clonal gametes are produced by a wide range of cytological mechanisms, from fully ameiotic (apomictic) processes to modified meiosis (automixis) [14,17]. In invertebrates and some cases of facultative parthenogenesis in vertebrates, unreduced gametes may form through suppression of polar body extrusion or gamete fusion [17–19]. In contrast, only two mechanisms have been observed in asexual hybrid vertebrates: premeiotic genome endoreplication and achiasmatic meiosis [5,17]. During endoreplication, germ cells duplicate their genomes prior to meiosis, allowing homologous chromosomes of the identical parental origin to pair and recombine normally [11,20–26]. In contrast, achiasmatic meiosis is characterized by the absence of homologous pairing and recombination; the reductional division is bypassed, while the equational division proceeds, resulting in unreduced clonal gametes [27–30].

One of the most widespread and evolutionarily successful asexual vertebrate complexes occurs in the genus *Carassius* [31–34]. This complex comprises sexual species (*C. auratus*, *C. gibelio, C. cuvieri, C. langsdorfii,* and *C*. *carassius*; ∼ 100 chromosomes), and the gynogenetic lineages (*C. gibelio* and *C. langsdorfii*; ∼ 150 chromosomes) [31,34–37]. The common ancestor of *Carassius* and *Cyprinus* underwent an allotetraploidization event approximately 14 million years ago [38,39]. Comparative chromosome-level assemblies show identical A and B subgenomes in both sexual and gynogenetic lineages, with paleotetraploid genomes (AaAbBaBb) and 100 chromosomes in *C. auratus/C. gibelio* (also referred to in the literature as “diploids”) and hexaploid genomes (AaAbAcBaBbBc) and 150 chromosomes in gynogenetic *C. gibelio* (often referred to in the literature as “triploids”) [40,41]. Hexaploid *C. gibelio* females are fertile and reproduce by gynogenesis, using sperm from related sexual species to activate egg development without genetic contribution [31,35,36]. Natural populations comprise not only numerous gynogenetic females and sexual species but also a small proportion of hexaploid *C. gibelio* males with decreased fertility (1.2–26.5%) [33,42].

Gametogenic alterations underlying gynogenesis in *C. gibelio* females remain complex and appear to vary among clonal lineages. In clone “D” of hexaploid *C. gibelio*, both bivalents and univalents were observed during meiotic prophase, along with the formation of first and second polar bodies [16]. These results suggested an additional genome duplication after fertilization of the egg by sperm from a different clone [16]. In contrast, clone “H” and laboratory-created hexaploid clones follow an ameiotic pathway characterized by the absence of synapsis and recombination, exclusive formation of univalents, and production of unreduced eggs. In this pathway, the first meiotic division is omitted via tripolar spindle formation, while the second division proceeds normally, yielding unreduced eggs and a single polar body [43,44]. Germ cell fusion has also been observed in laboratory hybrids of *C. auratus* × *Cyprinus carpio* [45].

Across asexual vertebrate complexes, reproduction is typically female-restricted, while hybrid males are sterile or show disrupted spermatogenesis, with only a few exceptions [11,22,26,46–49]. Notably, tetraploid/hexaploid *C. gibelio* represents a species in which both genetic (GSD) and environmental sex determination (ESD) occur [50,51,51–53]. A recent finding demonstrates that allotetraploid populations mostly rely on genetic factors, while triploid females produce males depending on rearing temperature [52]. Although hexaploid males are capable of producing sperm, their genetic contribution to offspring is rare, though their sperm can activate egg development [54]. Similarly, males from experimentally generated octoploid strains exhibit reduced sperm counts but retain fertility [30]. The presence of hexaploid males provides an opportunity to directly compare gametogenesis between sexes within the same genomic background. Our second aim is therefore to analyze meiosis in hexaploid *C. gibelio* males. As one of the few known asexual vertebrate lineages harbouring B chromosomes, *C. gibelio* represents a subject of particular interest. B chromosomes in asexual lineages are especially intriguing because, in the absence of canonical meiosis, the mechanisms maintaining or driving their transmission remain poorly understood. Another well-known vertebrate example is *Poecilia formosa*, where B chromosomes are of parental origin [55]. In *C. gibelio*, however, information regarding B chromosomes and their potential role in sex determination remains controversial. While their presence in hexaploid individuals has been clearly demonstrated [56,57], some studies suggest that specific B chromosomes may influence male determination in certain populations in China [51,58–60]. These findings have not been consistently replicated across populations, and the mechanisms underlying B chromosome inheritance in predominantly gynogenetic systems remain unclear.

In this study, we investigate gametogenesis in sexual tetraploid and asexual hexaploid *Carassius* lineages with three main aims: (1) to characterize meiotic chromosome behavior and determine whether asexual reproduction is associated with achiasmatic meiosis or alternative gametogenic modifications; (2) to evaluate sex-specific differences in gametogenesis, with particular emphasis on the reproductive potential of hybrid males; and (3) to assess the behavior and transmission of B chromosomes during meiosis in hexaploid asexual hybrids.

## Material and Methods

### Species and ploidy identification

In total, we investigated seven sexual tetraploid *Carassius gibelio* individuals (one juvenile female, one juvenile male, one adult male, and three adult females) (Supplementary Table S1). Further, we analyzed individuals obtained from 1) crosses of hexaploid *Carassius* females with hexaploid *Carassius* males (15 males and 5 females), 2) crosses of hexaploid *Carassius* females with *Scardinius erythrophthalmus* males (3 females), and 3) one wild-caught hexaploid female collected from the Olza River. The wild-caught individuals were identified via morphology and ploidy measurements by flow cytometry [61]. For these individuals, we examined mitotic and meiotic chromosomes, including synaptonemal complexes during pachytene (for males and females) and diplotene chromosomes (for females only). Gonadal microanatomy was additionally analyzed using confocal microscopy to assess the potential fertility.

### Laboratory crosses

All experimental procedures were conducted in accordance with Austrian law following the European Communities Directive 2010/63/EU on the protection of animals used for scientific purposes. Experimental fish originated from 16 wild-caught hexaploid *C. gibelio* from the Olza River system (Silesia, Czech Republic; [62]) and 20 tetraploid *C. gibelio* from Lake Neusiedl (Danube River system), provided by the Biological Research Station Illmitz.

Fish were held in 350 l aquaria in groups of 5-15 individuals depending on size. Light cycles and seasonal temperature were controlled following natural regimes. Water quality (temperature, oxygen content, levels of nitrate) was regularly monitored and adjusted via biweekly water refreshing. Fish were fed pellets and frozen chironomid larvae (blood worms) on a daily basis (except during winter time when food was provided every other day). During early May water temperature was increased from 12° to 16-17° within 3-4 days and artificial spawning substrate was provided (spawning brushes) to provoke natural spawning behavior. We refrained from using hormones to induce gamete ripeness. Typically, natural spawning occurred a few days later during early morning hours. When the beginning of spawning was observed, individual fish were caught with dip nets and gametes manually stripped by gently pressing the abdomen. Eggs of gynogenetic hexaploid *C. gibelio* were artificially inseminated with sperm from two different sperm donors, hexaploid males and *Scardinius erythrophthalmus*. Insemination with sperm from conspecific hexaploid males resulted in approximately 50% males and females, of which 12 males and 6 females were analyzed. Insemination of gynogenetic eggs with males of *Scardinius erythrophthalmus* resulted in 100% clonal female hexaploid offspring, of which three were analyzed. As controls, eggs of sexual tetraploid *C. gibelio* were inseminated with conspecific tetraploid *C. gibelio* males (4 females, 2 males analyzed). Fertilized or activated eggs were incubated at 20 °C in plastic trays to avoid temperature-induced masculinization due to elevated temperature [52,53]. Larvae hatched after five days and were subsequently reared in a closed water system (150 L). After seven months, juveniles were transferred to larger tanks in a flow-through system and raised to adulthood.

### Mitotic chromosome preparation

Mitotic metaphase chromosome spreads were prepared from the gills and kidneys of 21 individuals (3 tetraploid and 18 hexaploid) following standard protocols [63]. Adults were injected with 0.1% colchicine (10 µl/g body weight) for 3 hours; juveniles were kept in 0.05% colchicine solution for 4 hours. Metaphase chromosomes were stained with 5% Giemsa solution for 10 minutes at room temperature to initially assess chromosome number and morphology.

### Synaptonemal complex spread preparation and immunofluorescent staining

Synaptonemal complexes (SC) spreads during the pachytene stage of meiosis were prepared from testes of adult and juvenile males (2 tetraploid and 12 hexaploid) and ovaries of juvenile females (2 tetraploid and 9 hexaploid) following [64] and [65], respectively. Testis fragments were homogenized in 1× PBS, and 1 µl of the suspension was added to a 30 µl drop of hypotonic solution (1× PBS diluted 3 times in water) on SuperFrost® slides. After 20 minutes, cells were fixed with 2% paraformaldehyde (PFA) for 4 minutes and air-dried. Ovaries were homogenized in 0.2 M sucrose, and 20 µl of the suspension was dropped on slides, followed by the addition of 40 µl of 0.2 M sucrose and 40 µl of 0.2% Triton X-100 for 7 minutes. Cells were then fixed with 400 µl of 2% PFA for 16 minutes and air-dried. Slides were washed in 1× PBS and used for immunofluorescence.

SC lateral elements were visualized by rabbit anti-SYCP3 antibodies (ab15093, Abcam), and transverse elements were detected by chicken anti-SYCP1 antibodies. Anti-SYCP1 antibodies targeting the N-terminal region of zebrafish SYCP1 [66,67]. Recombination sites were detected using anti-MLH1 (ab14206, Abcam), and double-strand breaks were visualized using anti-Rad51 (GTX00721, GeneTex). After blocking in 1% BSA in 1× PBS with 0.01% Tween-20 for 20 minutes, slides were incubated with primary antibodies for 2 hours at room temperature. Afterward, slides were washed in 1× PBS and incubated for 1 hour with secondary antibodies: Alexa-488 goat anti-rabbit IgG (A-11008, Thermo Fisher Scientific), Alexa-594 donkey anti-chicken IgY (A-11042, Thermo Fisher Scientific), and Alexa-594 goat anti-mouse IgG (A-11005, Thermo Fisher Scientific). After washing in 1× PBS, slides were dehydrated in the ethanol series (50%, 70%, and 96%), air-dried, and mounted with Vectashield/DAPI (1.5 mg/ml, Vector).

### Diplotene chromosome preparation

Diplotene chromosomal spreads (so-called “lampbrush chromosomes”) were prepared from ovaries of three adult sexual tetraploid females and two asexual hexaploid females, according to an earlier published protocol [68] and further optimized for fish [69]. Ovaries were dissected and placed in the OR2 saline (82.5 mM NaCl, 2.5 mM KCl, 1 mM MgCl2, 1 mM CaCl2,1mM Na2HPO4, 5 mM HEPES (4-(2-hydroxyethyl)- 1-piperazineethanesulfonic acid); pH 7.4). Oocyte nuclei were isolated manually using jeweler forceps (Dumont) in the isolation medium “5:1” (83 mM KCl, 17 mM NaCl, 6.5 mM Na2HPO4, 3.5 mM KH2PO4, 1mM MgCl2, 1 mM DTT (dithiothreitol); pH 7.0–7.2) and immediately placed to one-fourth strength “5:1” medium with the addition of 0.1% paraformaldehyde and 0.01% 1M MgCl2. After removing the yolk, the oocytes’ nuclei were transferred to glass chambers attached to a slide filled with one-fourth-strength “5:1” medium. In glass chambers, the nuclear membrane was gently removed to release the nucleoplasm containing the chromosomes. Such a method ensures that each chamber contains chromosomal spread from a single individual oocyte. The slide was subsequently centrifuged for 20 min at 4 °C at 1500 g, fixed for 30 min in 2% paraformaldehyde in 1× PBS, and dehydrated in ethanol at 50%, 70%, and 96%. The glass chambers were removed before drying, and the diplotene chromosomes remained on the slides.

### Fluorescence *in situ* hybridization and fluorescent microscopy

Fluorescent *in situ* hybridization (FISH) was performed on metaphase spreads and lampbrush chromosomes from asexual species to detect B chromosomes using a PCR-labelled probe conjugated to biotin, according to CTAR3 [70]. For PCR, we used the following primers:

F 5’ TCCTCTTCTCACTCGGGAGCTG 3’

R 5’ ACTGGGTGATGGGACACTCTGG 3’

Furthermore, pancentromeric repeat probe (CTAR11, (Bishani et al. 2021) was labelled by PCR using the following primers:

F 5’ TCCTCTTCTCACTCGGGAGCTG 3’

R 5’ ACTGGGTGATGGGACACTCTGG 3’

Telomeric repeats were visualized using a probe prepared according to [71]. Metaphase slides were treated with 0.01% pepsin in 0.01 M HCl for 10 minutes, washed in 1× PBS, dehydrated through an ethanol series (50%, 70%, 96%), and air-dried. Lampbrush chromosome slides required no pretreatment. The hybridization mix contained 50% formamide, 10% dextran sulfate, 2× SSC, 5 ng/µl PCR-labeled probe, and 50ng/µl excess salmon sperm (ss)DNA. In the case of the telomeric probe, the hybridization mixture included 40% formamide, 10% dextran sulfate, 2× SSC, 10 ng/µl probe, and 50 ng/µl of tRNA. The probe was denatured at 86 °C for 6 minutes and then cooled on ice. Metaphase slides were denatured at 73 °C for 3 minutes, then dehydrated in cold ethanol (70%, 80%, 96%) and air-dried. The probe was applied to the slides, which were incubated overnight at room temperature in a humid chamber. Post-hybridization washes were performed three times in 0.2× SSC (2× SSC for the telomeric probe) at 44 °C for 5 minutes each. Streptavidin-Alexa 488 (Invitrogen) was used to detect biotin-dUTP; anti-digoxigenin antibodies (Invitrogen) were used to detect digoxigenin-dUTP. Chromosomal DNA was counterstained with Vectashield/DAPI (1.5 mg/ml, Vector).

Metaphase chromosomes were analyzed using a ZEISS Axio Imager.Z2 epifluorescence microscope with Olympus DP30BW and CoolCube 1 cameras (MetaSystems). Images were processed with IKAROS and ISIS, pseudocolored (blue: DAPI; red: anti-digoxigenin-rhodamine; green: streptavidin-FITC), and merged using Microimage. SC and diplotene spreads were examined with a Provis AX70 Olympus microscope using standard fluorescence filters. Images were captured with a DP30W CCD camera (Olympus) and processed with Olympus Acquisition Software. Final adjustments were performed using Adobe Photoshop version 26.

### Whole-mount immunofluorescence staining and confocal microscopy

Whole-mount immunofluorescent staining was performed on gonads from tetraploid (2 males and 2 females) and hexaploid (10 males and 6 females) individuals as described by [72]. Briefly, gonadal fragments were permeabilized in 0.5% Triton X-100 in 1× PBS for 4–5 hours at room temperature (RT), then washed in 1× PBS. Samples were incubated in 1% blocking reagent (Roche) for 1–2 hours before incubation with primary antibodies: mouse anti-α-tubulin (ab7291, Abcam) and rabbit anti-Vasa (DDX4, GTX116575, GeneTex). After overnight incubation at RT, tissues were washed and incubated overnight with Alexa-488-conjugated goat anti-rabbit IgG (A-11008, Thermo Fisher Scientific) and Alexa-594-conjugated goat anti-mouse IgG (A-11005, Thermo Fisher Scientific). Tissues were then washed and stained with DAPI (1 µg/µl, Sigma) overnight at RT.

Confocal imaging was performed using a Leica TCS SP5 mounted on an inverted Leica DMI 6000 CS microscope with a 63× HC PL APO objective. Fluorescence was excited with diode and argon lasers for DAPI, Alexa Fluor 488, and Alexa Fluor 594, and images were acquired and processed using LAS AF software (Leica Microsystems, Germany).

## Results

### Analysis of gonadal morphology of hybrid males and females

Gonadal microanatomy was analyzed using confocal microscopy. Germ cells were identified by Vasa staining, and developmental stages were distinguished based on nuclear morphology following established methods for *Cobitis*, *Hexagrammos*, and *Hypseleotris* species [72–74]. In tetraploid males and females, we revealed a normal distribution of germ cells, cells in meiosis and gametes (Figure 1A, C, E). In juvenile females, germ cells, pachytene and diplotene meiocytes were distributed throughout the ovary (Figure 1A). In males, we identified clusters of gonocytes, pachytene spermatocytes, metaphase I spermatocytes, and large groups of spermatids (Figure 1C). Meiotic cells displayed typical bipolar spindles with properly aligned bivalents at the metaphase plate (Figure 1E).

**Figure 1.**
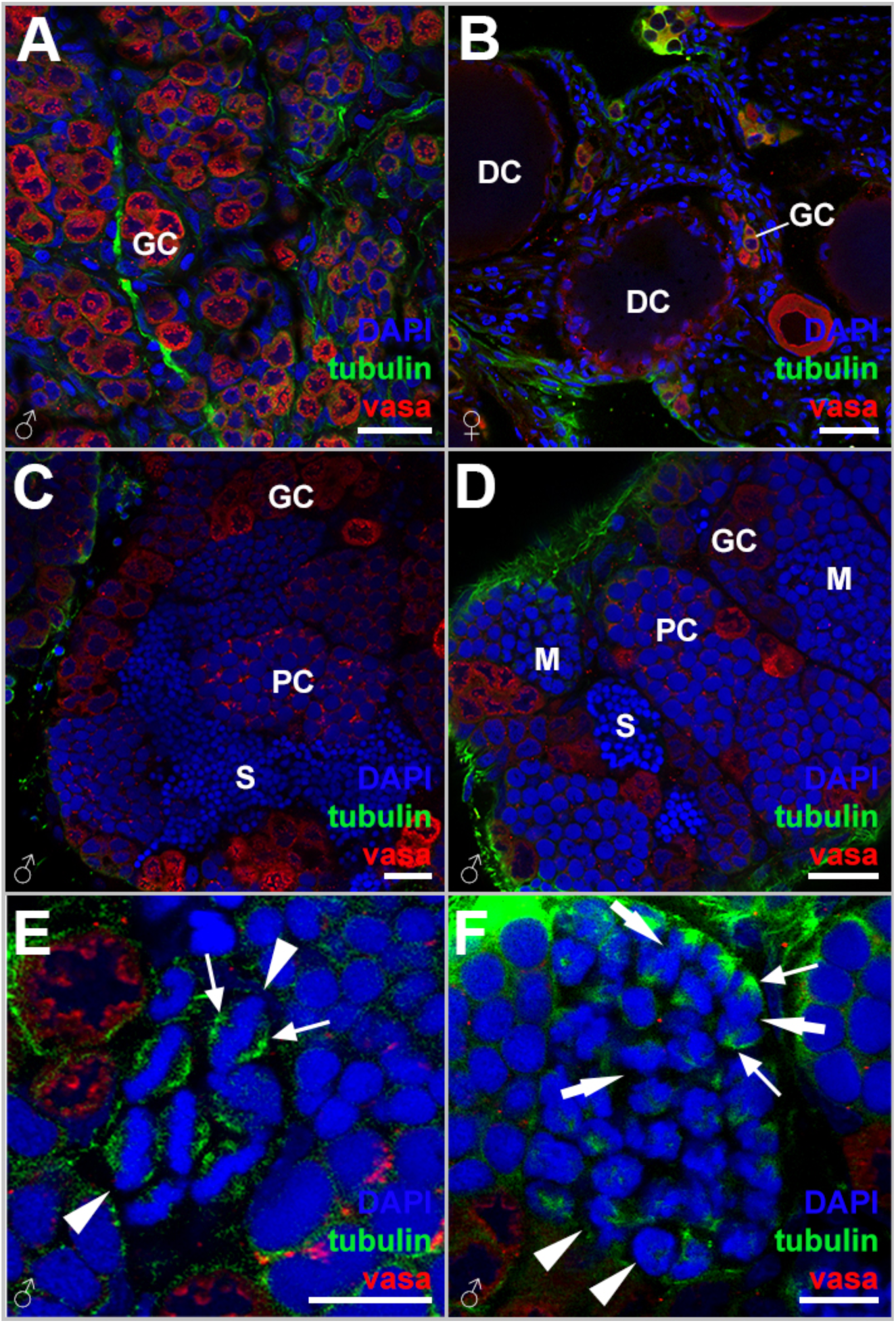
Gonadal microanatomy in tetraploid (A, C, E) and hexaploid (B, D, F) *Carassius* individuals. Single confocal sections of gonadal fragments and whole-mount immunofluorescent staining with antibodies against Vasa (red) and tubulin (green). DAPI visualizes chromatin (blue). Microanatomy of gonads from females (A, B) and males (C-F). Diplotene (DC) and gonial cells (GC) can be identified in gonadal sections of females (A, B). Cell types identified in the gonadal sections based on the morphology of males (C, D): S – spermatids, PC – cells in the pachytene stage of meiotic division, GC – gonial cells, M – meiocytes in meiotic division I. Meiotic divisions in tetraploid (E) and hexaploid (F) males. Thin arrows indicate bipolar spindle; arrowheads mark chromosomes in metaphases, thick arrows indicate chromosomes in anaphase. Scale bar (in A-D) = 50 µm; scale bar (in E, F) = 10 µm.

In hexaploid males, we also detected gonocytes, pachytene cells, and large clusters of cells during metaphase I, along with clusters of spermatids (Figure 1D). The analysis of cell clusters during meiotic divisions shows bipolar spindle formation; however, chromosomes failed to align at a defined metaphase plate and instead formed a diffuse chromosomal mass (Figure 1F). Despite this, the occurrence of anaphase-like figures indicates that cells proceed through division (Figure 1F). These observations suggest aberrant or modified meiotic divisions rather than canonical meiosis.

In hexaploid females, all stages of oogenesis were present, including pre-vitellogenic and vitellogenic oocytes, indicating retained reproductive capacity (Figure 1B).

### B chromosomes varied in size and copy number in asexual *Carassius* individuals

Mitotic analyses confirmed that somatic cells of sexual *C. gibelio* contain 4n = 100 chromosomes, consistent with previous reports [31,35,37,75], with no evidence of B chromosomes present. In contrast, somatic cells of hexaploid asexual *Carassius* exhibited approximately 6n = 150+Bs chromosomes, in agreement with earlier studies [31,34,35]. Using the CTAR 3 probe [70], which is known to produce very bright signals on B chromosomes, we detected hybridization signals along the entire length of B chromosomes, as well as occasional signals on pericentromeric heterochromatic regions of some autosomes (Figure 2; Supplementary Figure S1). In contrast to sexual individuals, in which no supernumerary elements were observed, B chromosomes were consistently present in all analyzed asexual individuals (n=17) (Supplementary Table S1). As previously reported for hexaploid *Carassius* individuals, B chromosomes showed substantial variability in both size and copy number [70]. They ranged from tiny microchromosomes to large macrochromosomes, with copy numbers ranging from 3 to 11 per individual (Figure 2; Supplementary Figure S1). This inter-individual variation was observed in both hexaploid males and females (Figure 2; Supplementary Figure S1). Two-color FISH using the B-specific tandem repeat (CTAR3, [70]) together with a pancentromeric repeat (CTAR11, [70]) demonstrated that both micro- and macro-B chromosomes lack the canonical pericentromeric repeat present on autosomes (Figure 2; Supplementary Figure S1).

**Figure 2.**
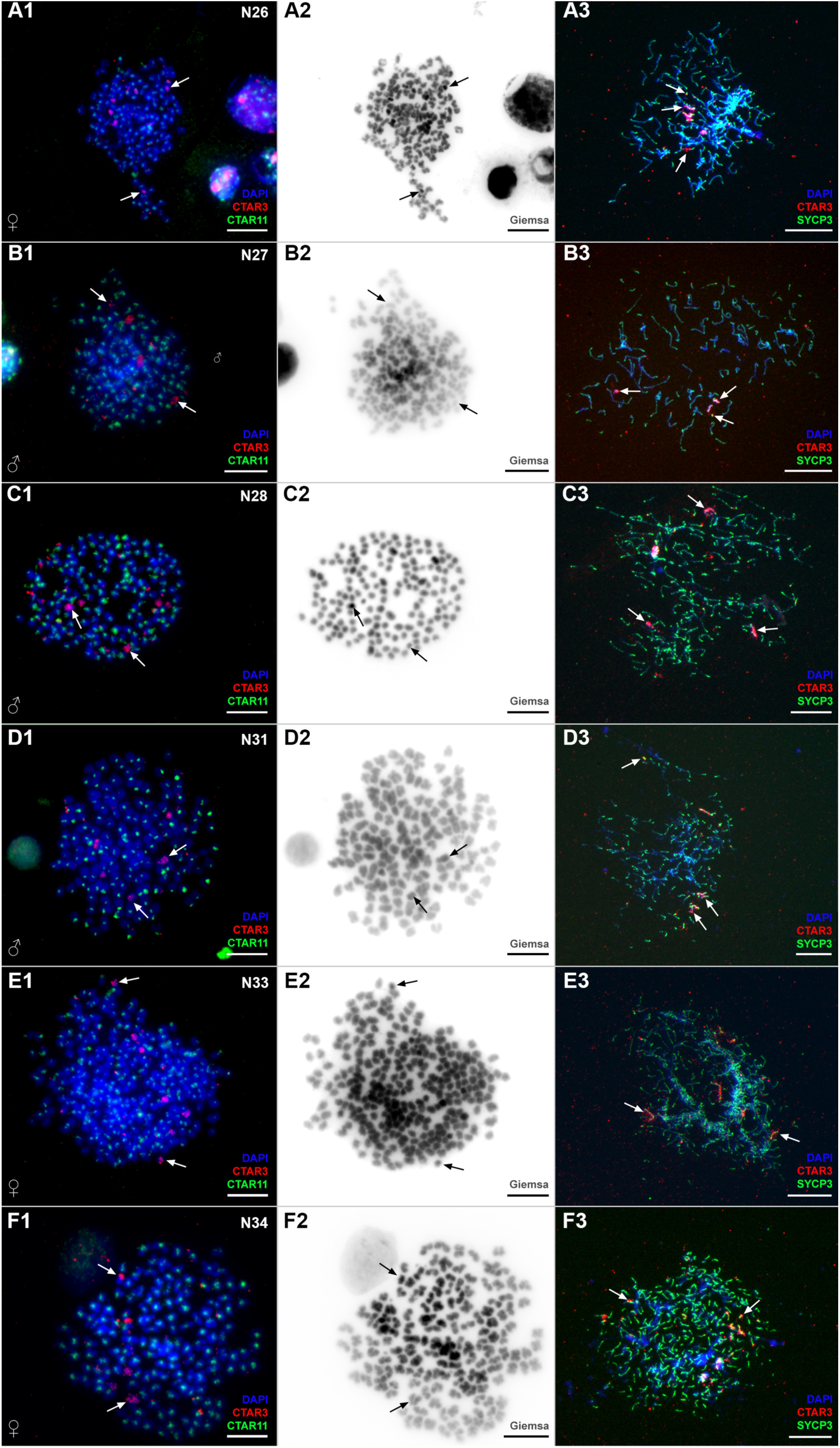
FISH-mapping of B chromosomes in mitosis and the pachytene stage of meiosis from the same individuals. Two-colored FISH with a probe detected intense signals on B chromosomes (red, CTAR3) and a probe to pancentromeric repeat (green, CTAR11) is localised on centromeres of all chromosomes except B chromosomes. Two-colored FISH is represented in A1, B1, C1, D1, E1, F1; corresponding Giemsa microphotographs are presented in A2, B2, C2, D2, E2, F2. Pachytene chromosome spreads of hexaploid individuals are represented in A3, B3, C3, D3, E3, F3. Lateral elements of synaptonemal complexes stained in green (SYCP3) and B chromosomes detected in red (CTAR3); chromatin is stained with DAPI, blue. Scale bar = 10 µm.

### Asexual *C. gibelio* males and females exhibit predominantly achiasmatic meiosis

We investigated pachytene chromosomes in two males and one female of tetraploid sexual individuals (Supplementary Table S1). To confirm proper bivalents formation during the pachytene stage of the sexual bioform, we stained synaptonemal complex components using SYCP1 (transversal element) and SYCP3 (lateral element). In sexual tetraploid individuals, pachytene analysis revealed 50 fully synapsed bivalents in both males and females, consistent with mitotic chromosome counts (Figure 3A, G). Each bivalent displayed at least one MLH1 focus, indicating normal crossing-over (Supplementary Figure S2A). Double-strand breaks, visualized by RAD51 staining, were most abundant during early pachytene and progressively declined during mid- and late pachytene (Supplementary Figure S2B), reflecting normal meiotic progression.

**Figure 3.**
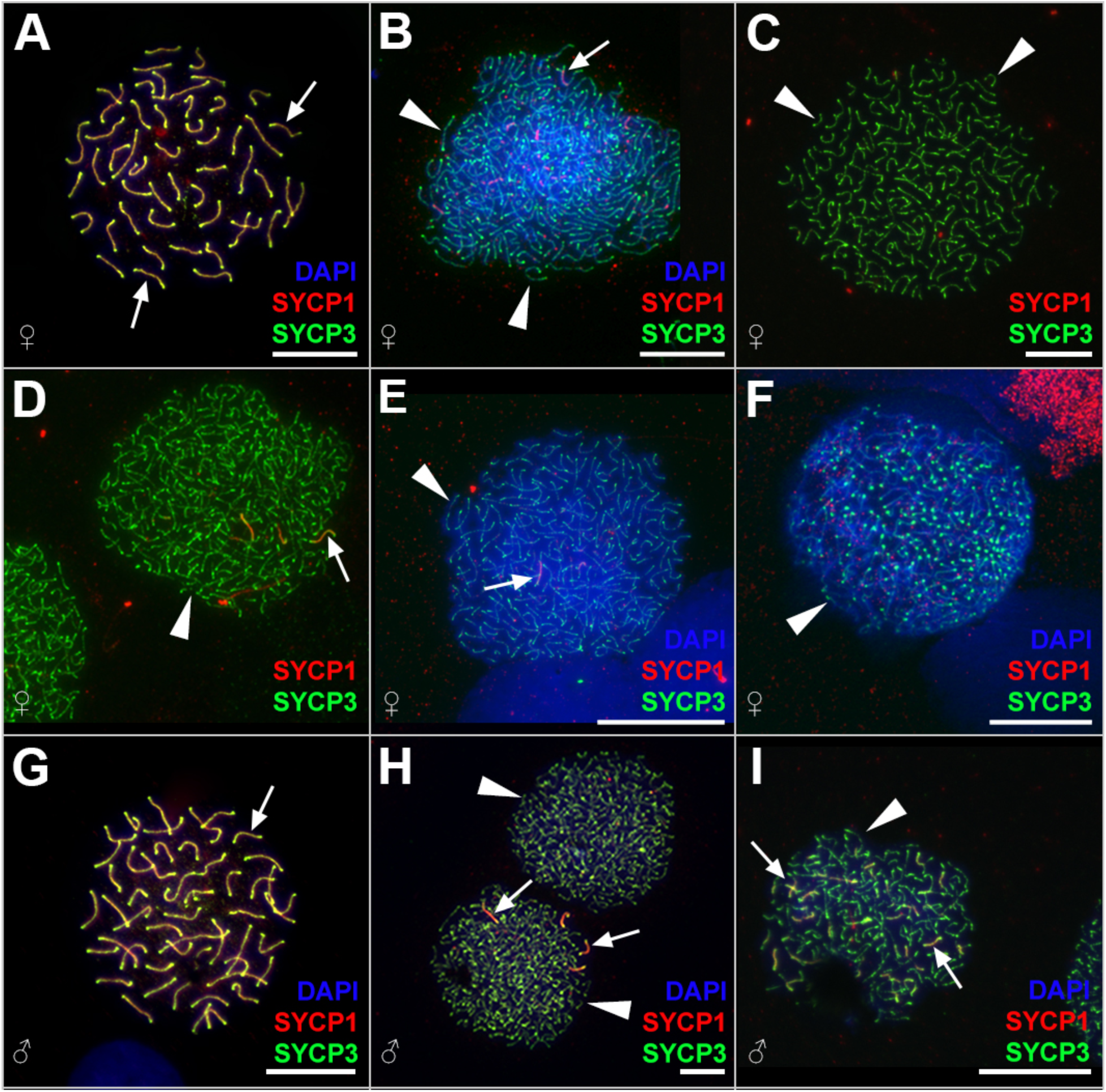
Pachytene chromosome spreads of tetraploid (A, G) and hexaploid (B-F, H, I) *Carassius* individuals. Staining for lateral (SYCP3) and transversal (SYCP1) components of synaptonemal complexes clearly shows the presence of 50 bivalents (indicated by arrows) in tetraploid females (A) and males (G). In hexaploid females (B-F) and males (H, I), around 150 univalents (indicated by arrowheads) have been found. In addition to meiocytes with exclusively univalents (C, F), meiocytes with occasional pairing (indicated by arrows in B, D, E, H) and intense pairing (I) have been observed. Arrows indicate bivalents and partially paired chromosomal fragments. Scale bar = 10 µm.

In contrast, analysis of hexaploid individuals (10 males, seven females) revealed extensive disruption of chromosome pairing. Among 1076 spermatocytes and 335 oocytes observed, the majority of cells (52% in males and 57 % in females) contained approximately 150 unpaired univalents with no sign of pairing (Figure 3C, F). In 488 spermatocytes (45%) and 146 oocytes (43%), we observed occasional pairing of 1-2 bivalents (Figure 3B-F, H), while the rest of chromosomes existed as univalents. Occasionally, in a minority of cells (3%) and exclusively in males, we detected spermatocytes showing more extensive pairing involving approximately 30 chromosomes, although complete synapsis was never observed (Figure 3I). Importantly, MLH1 foci were absent in all analyzed pachytene oocytes and spermatocytes of hexaploid individuals (Supplementary Figure S2C), indicating a lack of crossing-over even in those cells where occasional pairing occurred. RAD51 foci were also absent in pachytene oocytes (n = 28) and in most spermatocytes (n=81) (Supplementary Figure S2D-F). Only two spermatocytes exhibiting partial pairing showed chromosomes decorated with multiple RAD51 signals) (Supplementary Figure S2E).

Diplotene analysis supported these findings. In sexual tetraploid *C. gibelio* females, we observed 50 bivalents with clear chiasmata (Supplementary Figure S3). In contrast, asexual hexaploid females exhibited exclusively univalents (∼150 chromosomes) with no detectable chiasmata (Figure 4; Supplementary Figure S4). Although some univalents appeared transiently connected at telomeric regions, no chiasmata were detected, indicating the absence of stable pairing and crossing-over between homologous chromosomes (Supplementary Figure S4).

**Figure 4.**
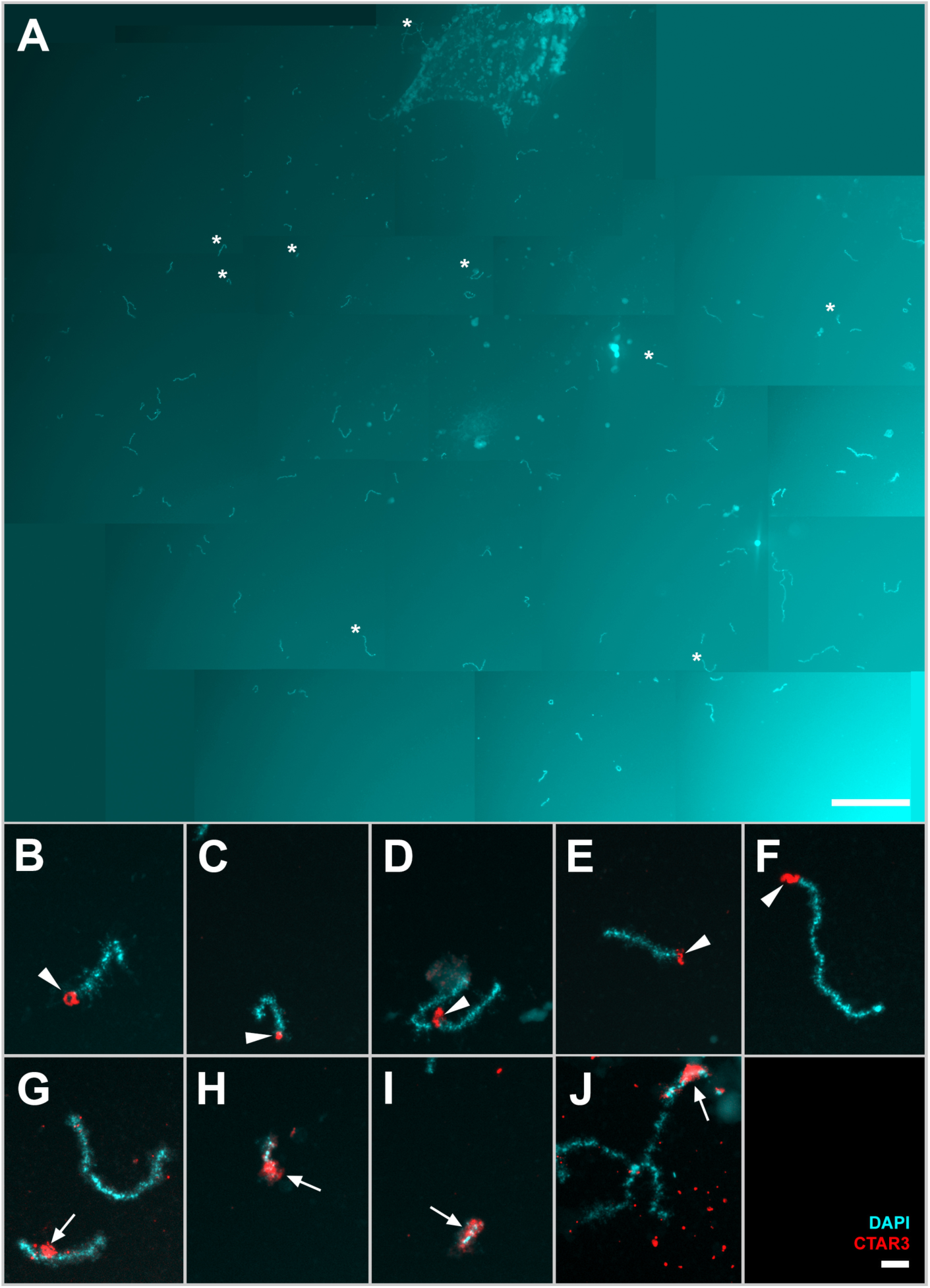
Diplotene chromosomal spread from an individual oocyte of a hexaploid *Carassius* individual. (A) Full chromosomal set with approximately 150 univalents. Since the chromosomal spread from the individual oocyte was large, 16 images were merged into one in the case of A. Asterisks indicate enlarged bivalents in A-D. Scale bar = 100 µm. FISH with a CTAR3 probe detected occasional hybridization signals on autosomes (B-F) and intense signals on B chromosomes (G-J). Scale bar = 10 µm.

### B chromosomes exist as univalents with no pairing during meiotic prophase in asexual lineages

Using a B-chromosome-specific probe, we visualized B chromosomes during pachytene and diplotene in eight hexaploid individuals (Figure 2, 4). At both stages, hybridization signals were observed along the entire length of the B chromosomes. During pachytene, B chromosomes behaved similar to the majority of other chromosomes in the asexuals and were consistently observed as univalents with no evidence of pairing (Figure 2, 3). During the diplotene stage, B chromosomes appeared as small univalents composed of several chromomeres and prominent lateral loops, indicating active transcription at this stage (Figure 4). Comparison of B-chromosome numbers in somatic cells and pachytene nuclei revealed similar copy numbers during meiotic prophase (Figure 2; Supplementary Table S1).

## Discussion

Asexual reproduction accompanied by polyploidization has been documented in several independent lineages of the genus *Carassius* that colonized Eurasia, partially due to human activity. Despite extensive research, the differences and discrepancies among lineages and forms require detailed analysis of gametogenesis to elucidate the evolutionary success of this group. Here, we demonstrated that hexaploid asexual *C. gibelio* lineages undergo profoundly altered meiosis characterized by the absence of homologous pairing, recombination, and chiasma formation. By comparing male and female meiosis, we found that both sexes initiate achiasmatic meiosis during prophase. However, females are fully fertile, whereas males exhibit reduced fertility, likely due to their inability to alter the meiotic division. In addition, we documented extensive variability of B chromosomes and described their behavior during meiosis of asexual lineages.

### Both males and females from hexaploid *C. gibelio* exhibit achiasmatic meiosis

We found that in both sexes, most meiocytes lacked synapsis where chromosomes progressed through the pachytene and diplotene stages of meiosis exclusively as univalents. The absence of crossing-over and double-strand break foci further indicates a failure to establish canonical meiotic recombination and chromosomal synapsis. These observations are consistent with previous cytological studies of hexaploid *Carassius* clones [30], which also reported extensive asynapsis and a lack of recombination.

Notably, in addition to fully asynapsed chromosomes, we and others [30] observed a small subset of oocytes and spermatocytes with partial synapsis, accompanied by intense double-strand break formation. Such cells are unlikely to complete meiosis successfully and probably represent sporadic attempts to initiate canonical recombination rather than an alternative reproductive pathway. This contrasts with the gynogenetic fish with achiasmatic meiosis *Poecilia formosa*, in which only fully asynaptic oocytes have been described [29]. The presence of two distinct oocyte populations has previously been observed in hybrid lineages of fish and lizards that employ premeiotic genome endoreplication, highlighting the complexity of gametogenic alterations towards asexuality [24,72,76–78]. In such lineages, only 2–5% of oocytes successfully duplicate their genomes and produce diploid gametes through normal meiosis, while the remaining oocytes retain the original ploidy level and arrest during pachytene. Interestingly, the proportion of such “wasted” oocytes appears to be considerably lower in hexaploid *Carassius*. Nevertheless, to determine the exact proportion of oocytes capable of completing subsequent meiotic divisions in *C. gibelio* needs further investigation.

Achiasmatic meiosis has been documented in only a limited number of asexual vertebrate lineages, namely *Poecilia formosa* [27,29] and clone H of *C. gibelio* (our data and [30]), indicating that complete suppression of pairing and recombination accompanied by altered meiotic divisions represents a comparatively rare route to asexuality in hybrid vertebrates. In contrast, premeiotic genome endoreplication is far more widespread among asexual hybrid vertebrates, and has been reported across diverse taxonomic groups [11,20,23–26,79–81]. Nevertheless, since both achiasmatic meiosis and premeiotic endoreplication have been so far documented only in hybrid strains of vertebrates, it is likely that both programs require fundamental alterations of the canonical gametogenic program, triggered in hybrid genomic backgrounds. In this context, it is worth mentioning that premeiotic genome endoreplication is far more common among hybrid asexuals, than achiasmatic meiosis. In addition to *P. formosa* [27,29] and *C. gibelio* (our data; [30]), and it is quite likely that *C. langsdorfii* follows achiasmatic meiosis, since Yamashita and co-authors ([28] observed skip page of the first meiotic division. Interestingly, however, at least in some spermatocytes, chiasmata as well as bi- and trivalent formations were observed among certain chromosomes during meiotic prophase in *C. langsdorfii* [82]. To add even more complexity, we recently noted similar achiasmatic-like meiotic phenotypes, characterized by the absence of double-strand break formation, homologous pairing, and crossovers, in hybrids of catfish *Clarias macrocephalus* × *C. gariepinus* [78] where, premeiotic genome endoreplication was otherwise detected in majority of cells. This suggests that both these alternative mechanisms may be involved in parallel [78]. Together, these observations suggest that achiasmatic meiosis and premeiotic endoreplication may represent alternative outcomes of hybrid-induced disruption of meiotic regulation. Whether these two aberrant pathways share common mechanistic triggers or represent distinct evolutionary solutions to similar constraints remains an open question.

### Sex-specific differences and fertility consequences in hexaploid *C. gibelio* males and females

In *C. gibelio*, achiasmatic meiosis was observed in both hexaploid females and males exhibited. This pattern contrasts with most gynogenetic lineages that rely on premeiotic genome endoreplication, whereby males are unable to duplicate their genomes and consequently remain sterile or show strongly reduced fertility due to aberrant chromosome pairing [11,26,49,83]. Even in triploid males derived from *P. formosa* clonal females, synapsis is present on some bivalents, suggesting the different gametogenic regulations compared to *P. formosa* females [84]. Despite sharing a similar achiasmatic meiotic program, reproductive outcomes in hexaploid *C. gibelio* differ between the sexes. Females completed oogenesis, including the formation of vitellogenic oocytes, indicating functional fertility. In contrast, males displayed extensive meiotic abnormalities, limited progression through spermatogenesis, and reduced gonadal development, consistent with partial or complete sterility. Nevertheless, rare sperm produced by hexaploid males can activate egg development without contributing genetically (our data, [54]). Similarly, a rare sperm which can trigger gynogenetic reproduction was reported for *P. formosa* males [85].

These pronounced sex-specific differences in fertility likely arise from differential abilities to bypass or modify meiotic divisions. In hexaploid *C. gibelio* and *C. langsdorfii,* the reductional meiotic division fails due to aberrant spindle organization, most commonly through tripolar spindle formation [28,43,44]. These meiotic alterations, occurred in females, prevents proper attachment and segregation of univalent chromosomes during first meiotic division. The second meiotic division proceeds normally via bipolar spindle formation, leading to chromatid separation and the formation of only one polar body and an egg with genome composition identical to that of the maternal somatic cells [28,43,44]. Notably, despite these profound cytological deviations, key meiotic cell-cycle regulators, including maturation-promoting complex and cyclins, appear broadly comparable to those in sexual species [28,43,44].

In males, however, we consistently observed bipolar spindle formation without evidence of tripolar spindles during meiosis (Figure 1D). Chromosomes failed to establish proper attachments to the spindle, suggesting that the reductional division is not effectively bypassed. Nevertheless, the presence of anaphase of meiotic division suggests the possibility of chromosome segregation at least in some meiocytes. Consequently, although both sexes share similar premeiotic and early meiotic disruptions, only females successfully resolve these constraints to produce fertile gametes. These differences in spindle organization and meiotic progression provide a mechanistic explanation for the striking divergence in fertility between males and females in asexual *Carassius* lineages.

### B chromosomes in asexual *Carassius*

A striking feature of asexual *C. gibelio* lineages is the consistent presence of B chromosomes. For a long time, high diploid numbers and limited cytogenetic investigations have underscored the presence of B chromosomes in this species. Thus, in many early publications, these elements have been described as microchromosomes only. We observed pronounced interindividual variation in both the copy number and size of B chromosomes, as well as in their molecular content. Specifically, the number of B chromosomes ranged from 3 to 11 per individual, and their size varied from microchromosomes to medium-sized chromosomes. Comparable polymorphism in B-chromosome number and morphology has been reported in east asian, Polish, and Siberian populations of *C. gibelio* [58,70,86]. Despite this variability, some reports indicate that B chromosomes share highly similar sequence composition, suggesting a common evolutionary origin [34,58]. In some East Asian populations, it was discovered that there is a subset of micro-B chromosomes specific to males, which generally carry a higher number of additional chromosomes, and these elements were suggested to play a role in sex determination [58–60]. However, we found no evidence for sex-biased distribution of B chromosomes in our samples, consistent with recent observations [52].

Notably, a pancentromeric repetitive sequence was detected on all autosomes but was absent from B chromosomes. This observation suggests that the repeat is either absent from B chromosomes, present at copy numbers below the FISH detection threshold, or sufficiently diverged to prevent probe hybridization. A similar divergence in pericentromeric repeat composition has been reported for germline-restricted chromosomes in nightingale species and fungus gnat, where distinct centromeric or pericentromeric sequences may be associated with their specific transmission dynamics [87,88]. We further demonstrated that B chromosomes behave as univalents during meiosis and persist through pachytene and diplotene stages. Importantly, their copy number remains consistent between somatic and meiotic cells, indicating stable transmission without elimination during ontogenesis.

In sexual species, B chromosomes typically face challenges in segregation and propagation, as successful transmission often requires either bivalent formation or specialized mechanisms that bias chromatid segregation or meiotic disjunction. As univalents, supernumerary chromosomes are commonly prone to loss during meiosis in sexually reproducing individuals [89,90]. In contrast, in asexually reproducing *Carassius* lineages, achiasmatic meiosis may facilitate the stable inheritance of B chromosomes by allowing them to behave similarly to autosomes during gametogenesis, thereby promoting their persistence and accumulation.

## Acknowledgements

We would like to thank Thomas Zechmeister, Richard Haider and Rudolf Schalli from the Biological Station Illmitzl for fish from Lake Neusiedl, and Lukas Kalous for fish from Silesia. The current work was performed in the framework of the Research Focus “Alpine Space – Man & Environment” University of Innsbruck, Austria,

## Funding

This study was supported by the Czech Academy of Sciences (RVO: 67985904 to D.D. K.J.), the Czech Science Foundation (GA CR) (23-07028K to D.D, K.J., H.Z., and K.K.). V.T. was supported by the Marie Skłodowska-Curie Actions - COFUND project, which is co-funded by the European Union (MERIT - Grant Agreement No. 101081195).

## Conflicts of Interest

The authors declare no conflict of interest.

## Data Availability

The data underlying this article are available in the article and in its online supplementary materials.

## Author contribution

DD: Conceptualization; Funding acquisition; Investigation; Supervision; Writing—original draft; Writing—review & editing. DKL: conceptualization; provision of fish material, animal husbandry, artificial crossing; manuscript writing; HZ, KK, VT: chromosome preparation, FISH experiments, manuscript writing. JW: artificial crossing; animal husbandry. MS: conceptualization; YI: antibodies preparation, manuscript review & editing; all authors: manuscript review & editing.

## Supplementary Materials

**Figure S1.**
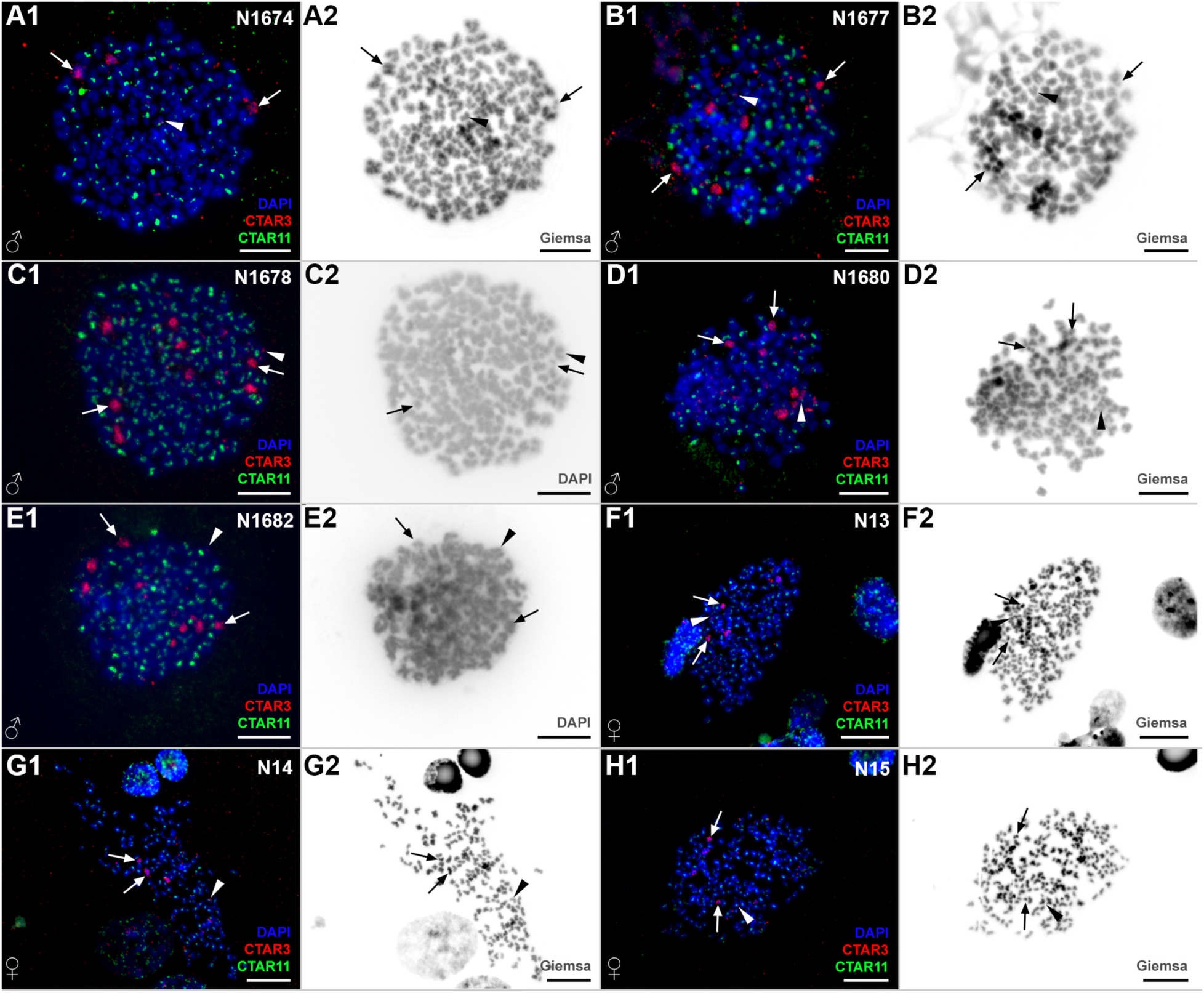
Two-colored FISH of B chromosome-specific probe and pancentromeric repeat in mitosis of hexaploid males (A1-E2) and females (F1-H2) *Carassius* individuals. FISH with a CTAR3 probe detected occasional hybridization signals on autosomes (red, shown by arrowheads) and intense signals on B chromosomes (red, shown by arrows). Pancentromeric repeat (gree, CTAR11) is localized on centromeres of all chromosomes except B chromosomes. Two colored FISH microphotographs are represented in A1, B1, C1, D1, E1, F1, G1, and H1. Corresponding Giemsa microphotographs are presented in A2, B2, D2, F2, G2, H2, and DAPI in grayscale is presented in C2 and E2. Scale bar = 10 µm.

**Figure S2.**
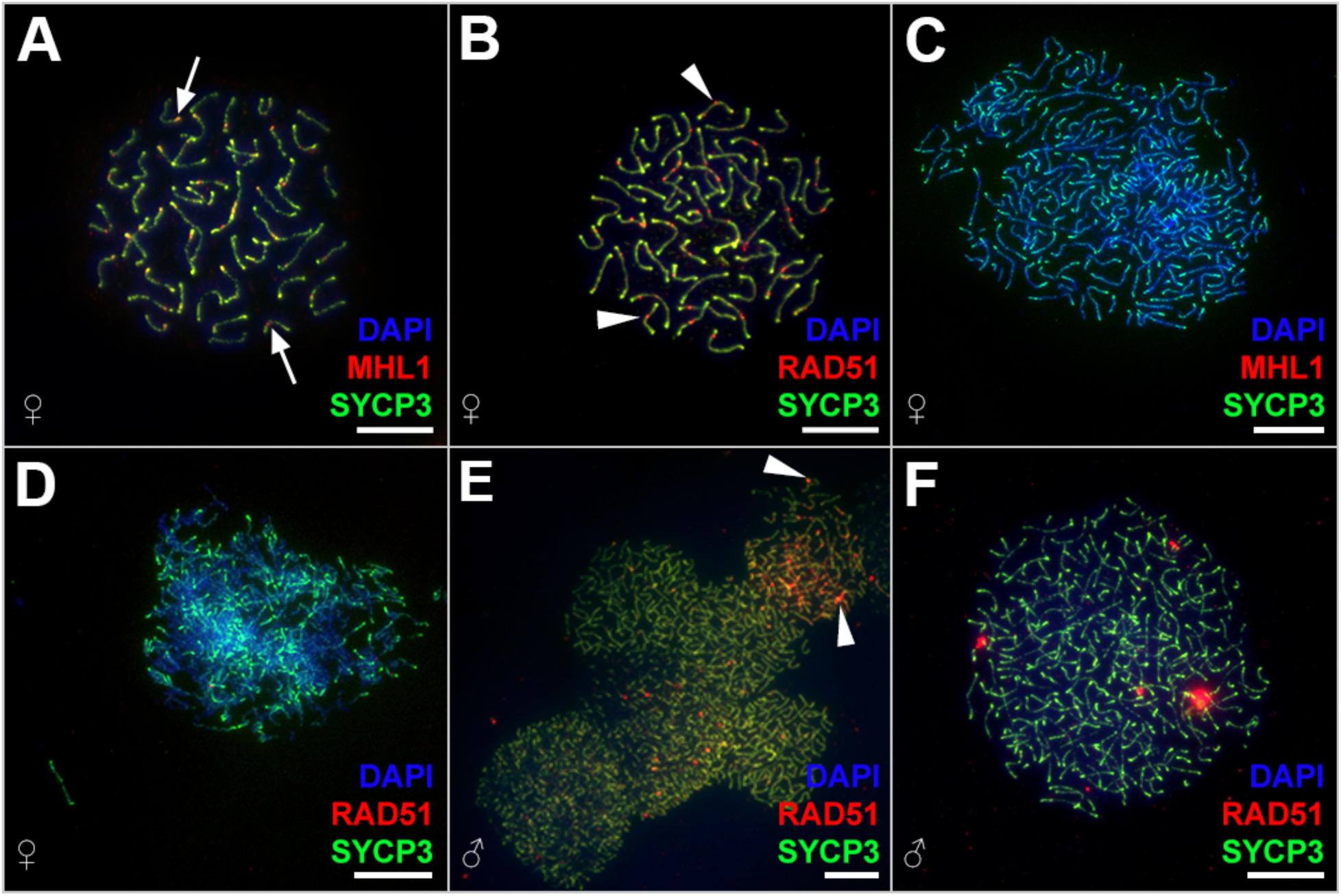
Pachytene chromosome spreads of tetraploid (A, B) and hexaploid (C-F) *Carassius* individuals. MLH1 staining visualizes crossover loci (pointed by arrows) in the sexual tetraploid female (A), while no crossovers were detected in oocytes from the hexaploid female (C). RAD51 staining (indicated by arrowheads) shows the presence of DSBs formation in tetraploid bivalents (B) but not the majority of spermatocytes (E, F) and oocytes (D) from hexaploid individuals. Nevertheless, DSBs were found in occasional spermatocytes of hexaploid males (E). Scale bar = 10 µm.

**Figure S3.**
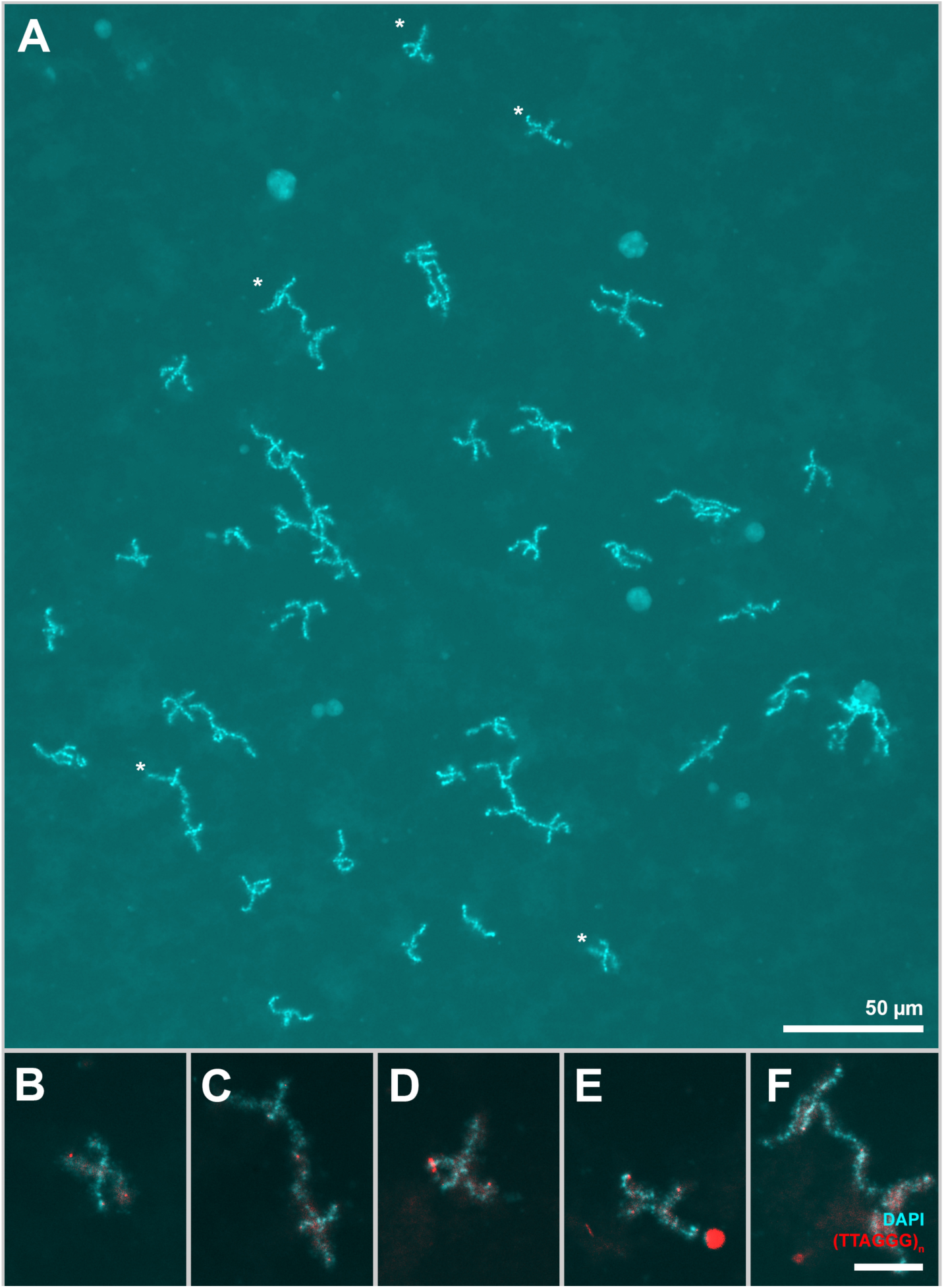
Diplotene chromosome spreads from an individual oocyte of a tetraploid *Carassius* individual. (A) Full chromosomal set with approximately 50 bivalents. Since the chromosomal spread from the individual oocyte was large, two images were merged into one in the case of A. Asterisks indicate enlarged bivalents in B-F. Scale bar in A = 50 µm. FISH with a (TTAGGG)n probe detected ends of individual chromosomes united in bivalents (B-F). Scale bar = 10 µm.

**Figure S4.**
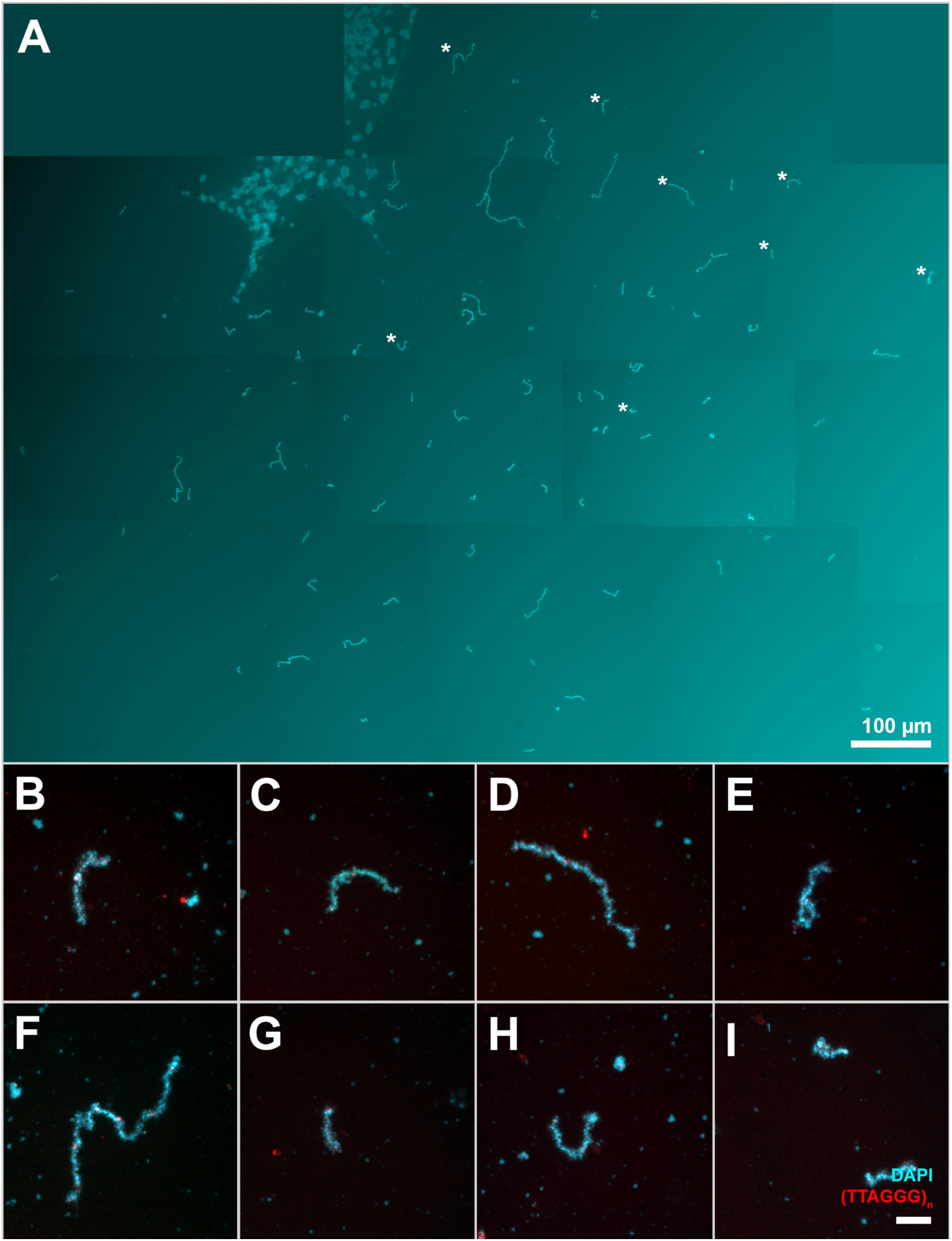
Diplotene chromosome spreads from an individual oocyte of a hexaploid *Carassius* individual. (A) Full chromosomal set with approximately 150 univalents. Since the chromosomal spread from the individual oocyte was large, 15 images were merged into one in the case of A. Asterisks indicate enlarged bivalents in B-I. Scale bar in A = 100 µm. FISH with a (TTAGGG)n probe detected ends of individual chromosomes existing as univalents (B-I). Scale bar = 10 µm.

**Supplementary Table S1. The list of individuals and methods used in the study.**

## Notes

### Competing Interest Statement

The authors have declared no competing interest.

